# A custom two-in-one HIST and line-scanning confocal excitation module

**DOI:** 10.64898/2026.09.14.751507

**Authors:** Christian Arthur, Daniel E. Milkie, Andrew J. Ulmer, Justin P. Ellis, Hark Kyun Kim, Feifei Song, Alejandro Chavez, Wesley R. Legant

## Abstract

Fluorescence microscopy applications often require specialized instruments that are optimized for different experimental goals. Here, we present a reconfigurable microscopy module that integrates highly inclined swept tile (HIST) illumination for high-sensitivity single-molecule imaging and line-scanning confocal microscopy for rapid and optically sectioned volumetric acquisition. The system shares major hardware components, including lasers, scanning optics, and detection hardware, while employing unique beam shaping pathways to enable rapid switching between modalities without realignment. We characterize the module performance by measuring the excitation beam profiles, the point spread functions (PSF), and the optical transfer functions (OTF) across 40x, 60x, and 100x magnifications and demonstrate imaging applications including diffraction-limited fixed and live-cell volumetric imaging, fluorescence recovery after photobleaching, and super-resolution DNA-PAINT and single particle tracking (SPT). We also demonstrate the capability to execute multimodal imaging workflows by performing confocal imaging for chromatin density classification correlated with SPT data of nuclear proteins with diverse functions. Together, these results demonstrate a versatile imaging platform capable of supporting complementary fluorescence imaging modalities within a single instrument.

## 1 Introduction

Due to its high sensitivity, low background, and protein specificity, fluorescence microscopy has become an indispensable technique for biological research. Modern fluorescence microscopes come in a variety of fixed configurations that choose among the trade-offs between optical sectioning, field of view (FOV), and acquisition speed^1^. As a result, different experimental needs often require separate, specialized instruments with unique hardware that increases cost and limits experimental flexibility. For example, single-molecule imaging studies are commonly performed using widefield, total internal reflection fluorescence (TIRF), or highly inclined and laminated optical sheets (HILO)^2–6^ combined with camera-based detectors that maximize photon collection efficiency and throughput. In contrast, volumetric imaging often utilizes confocal microscopes with pinholes and point detectors to reject out-of-focus emission and maximize optical sectioning^7,8^. However, over the past decade, advances in sCMOS cameras with rolling shutter readout have enabled the creation of new imaging configurations with the potential to address both use cases in a single instrument.

Single-molecule imaging can be performed at increased sample depths while still maintaining high signal-to-noise by utilizing HILO. To generate a HILO beam, excitation light is focused to a point on the objective pupil just inside what would be needed for TIRF^9^. This method produces an oblique beam that passes through the objective focal plane at the center of the field of view (FOV). However, because the beam is angled rather than coincident with the detection plane, the effective field-of-view depends on the illumination angle and the beam thickness, limiting the original demonstration to FOV ranging between 15 and 40 µm and sheets with thicknesses between 6 and 10 µm, respectively^9^. More recently, highly inclined swept tile (HIST)^10^ and oblique line scan (OLS)^11,12^ microscopy have overcome the trade-off between illumination thickness and FOV by sweeping an elongated, tile-like beam and synchronizing it with a rolling shutter detection on an sCMOS camera. This combination enables rapid imaging over large areas with a high signal-to-background ratio (SBR) and is well suited for single-molecule imaging and single-molecule in situ hybridization studies. However, to attain a per-line integration time that is sufficient to capture the limited photons from single-molecule emission, HIST and OLS utilize fairly thick oblique sheets (3 µm) that are coupled to large integration widths for the rolling shutter (∼84 camera pixels corresponding to approximately 9 µm at the sample plane or roughly 15 Airy units with the high numerical aperture lenses used in these studies). As such, these methods optimize photon collection over optical sectioning and axial resolution, thus limiting suitability for large-volume and deep tissue biological samples.

Fortunately, line-scanning confocal microscopy can be implemented with the same hardware by utilizing different optical components upstream of the scanning galvo^13^. To generate a line-scanning confocal beam, the light illuminates a centered and focused line spanning the full diameter of the objective pupil. This approach produces a diffraction-limited line of illumination that propagates along the objective optical axis with a focus that coincides with the detection plane. When coupled with a narrow rolling shutter integration width of approximately 1 Airy unit, this method enables real-time rejection of out-of-focus and scattered light, improving contrast and SBR while maintaining high imaging speed.

These two approaches offer complementary strengths but are usually implemented in separate instruments, leading to duplication of costly components such as laser sources and detection hardware, as well as limiting the ability to perform multimodal experiments within a single imaging workflow. Here, we present a custom microscopy solution that integrates HIST and line-scanning confocal within a single, reconfigurable instrument. The system incorporates two excitation modules that can be selectively engaged, allowing rapid switching between imaging modes. In this way, the platform effectively functions as two microscopes at different times, enabling both large-FOV, high-SBR inclined illumination imaging and confocal line-scanning with enhanced optical sectioning. By reusing key components, such as laser sources, galvo mirror, and scan lenses, the system reduces overall cost while expanding its experimental capability. This design also provides the ability to automatically switch between imaging modes, enabling seamless complex imaging experiments with enhanced efficiency. After characterizing the instrument performance for each modality across magnifications ranging from 40x – 100x, we demonstrate several use cases across both live and fixed samples. We also provide a detailed parts list and computer-assisted design (CAD) models for those wishing to replicate the system.

## 2 Results

### 2.1 Custom module design and concept

#### 2.1.1 Mechanical design

The custom imaging system incorporates both the line-scanning confocal and HIST modalities in a single module (**Figure 1A**) that can be integrated with a commercial microscope body (Nikon Ti2-E) and with a commercial API that can be accessed from a variety of external software packages. This allows for seamless integration with existing motorized stages, focus control, fluorescence filter sets, and environmental control accessories. We maximized the hardware shared between the two imaging modalities to reduce system complexity and overall cost. Consequently, the laser sources, camera, galvo mirror, scan lens, tube lens, and control hardware, which represent roughly 80% of the total instrument cost (**Supplementary Table 1**), are shared between both modes.

**Figure 1.**
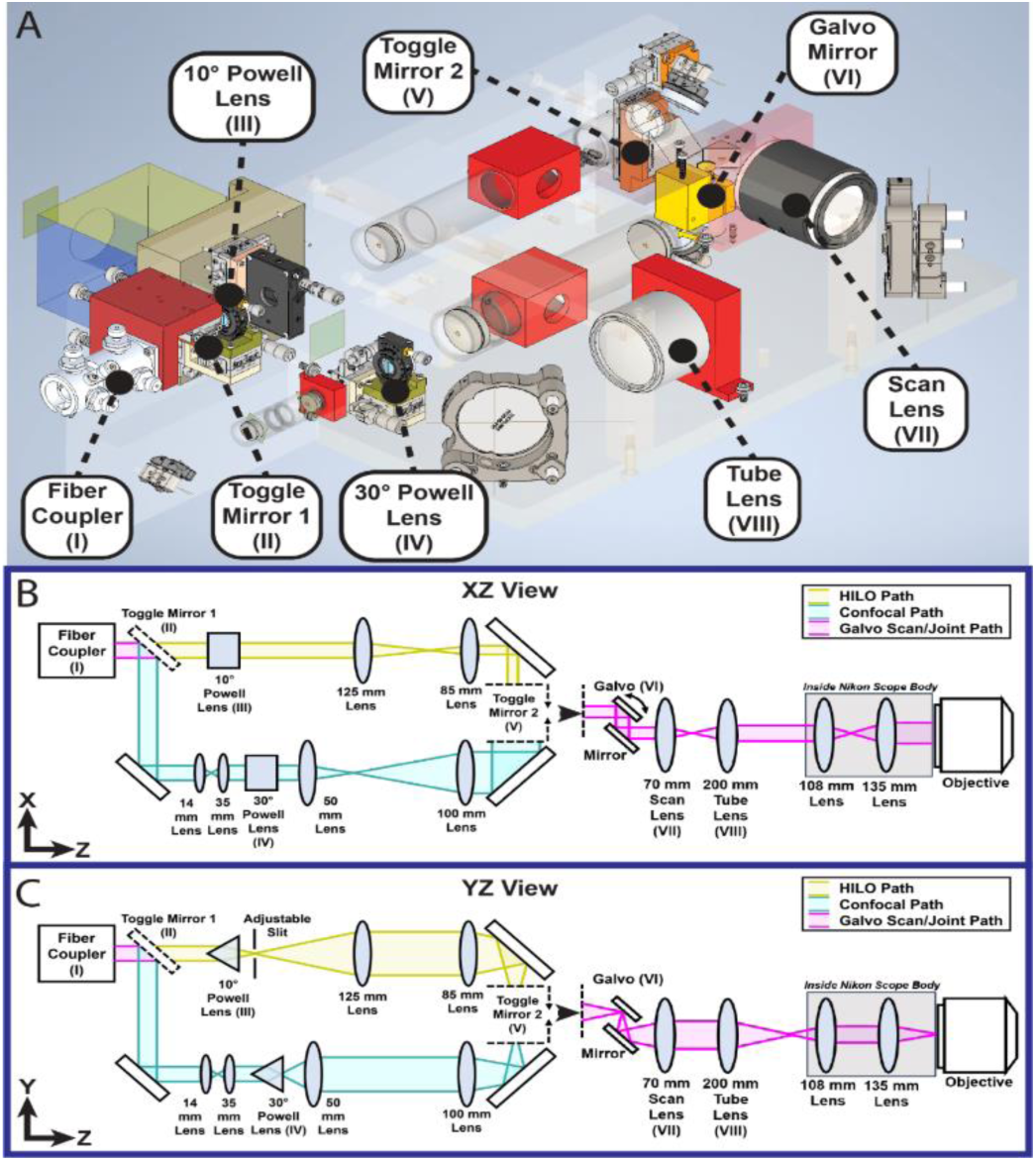
Custom HIST-Line scanning confocal excitation module design. **(A)** 3D model of the custom excitation module. **(B)** XZ view of the optical path of the custom excitation module. **(C)** YZ view of the optical path of the custom excitation module. The diagrams’ coordinate system follows the sample coordinate system.

Because each mode requires different beam profiles at the sample, the input beam from the optical fiber is picked off by a toggle mirror and diverted into one of two separate beam-shaping pathways, each optimized for either line-scanning confocal or HIST illumination. These separate paths are then directed onto a single scanning galvo utilizing a knife-edge mirror and periscope before being projected through a shared scan lens and tube lens (**Figure 1B, C**). To optimize mechanical stability, spatial footprint, and assembly, we designed a custom-machined frame that integrates adjustable kinematic mounts at key locations in the optical path. This approach reduced the overall module size to roughly 55 x 25 x 15 cm while still giving sufficient adjustment for precise optical alignment.

#### 2.1.2 Optical design

We designed the confocal system to generate a diffraction-limited line at the sample for three different objective magnifications and numerical apertures (NA): 40x /1.25NA, 60x/1.27NA, and 100x/1.35NA. All configurations utilized a 15 mm square sCMOS camera chip. In these scenarios, the 40x/1.25NA lens requires the largest beam size and scan angles. Thus, we designed an optical path to focus the light into a ∼18 µm wide full width at half maximum (FWHM) line that spans the entire 12.5 mm diameter of the 40x objective pupil while accounting for the additional relay optics that are internal to the microscope body. When operating at other magnifications and NA, rather than introducing additional beam shaping optics, we instead overfill the objective pupil, favoring reduced cost and simplicity over excitation efficiency. With this, we calculated the angles and magnifications between the pupil-conjugate scanning galvo and objective pupil necessary to cover a camera-limited 375 µm field of view. To generate a uniform line profile at the pupil, we designed a beam shaping path that first expands the beam 2.5x from the fiber, converts the collimated Gaussian beam into a fan distribution utilizing a 30° Powell lens, and then magnifies this 2x onto the scanning galvo using 4F relay optics (**Figure 1B, C**).

The HIST imaging path shares the same scanning galvo and downstream optics as the confocal path but utilizes a separate beam shaping path with no initial beam expansion and a 10° Powell lens followed by 0.68x demagnifying relay optics. While the HIST beam width at the sample is fixed by the Powell lens and relay optics, the beam angle and thickness can be tuned via a mechanical slit that restricts the angular spectrum of rays subtended by the HIST illumination and by controlling the location of a HIST-path-specific mirror on a translation stage that shifts the center of this spectrum to different locations in the pupil.

Following the selection of optical elements, the path from the scanning galvo to the objective pupil, assuming a perfect objective lens, was simulated with a commercially available ray-tracing software (Zemax OpticStudio and Code V) to guide the precise placement of components and optimize beam propagation quality throughout the system. The resulting spot diagrams demonstrated that beam quality was maintained throughout the scan range with minimal degradation at the field edges (**Supplementary Figure 1**), confirming that the optical design can support the intended imaging area without introducing significant scan-dependent aberrations. However, as noted in more detail below, because the beam cannot be fully recollimated after the Powell lenses, we were unable to achieve uniform illumination over the full design spec along the Y-axis (**Figure 2A, B)**. This effect was worse for the confocal imaging mode due to its larger-angle Powell lens (**Supplementary Table 2**).

**Figure 2.**
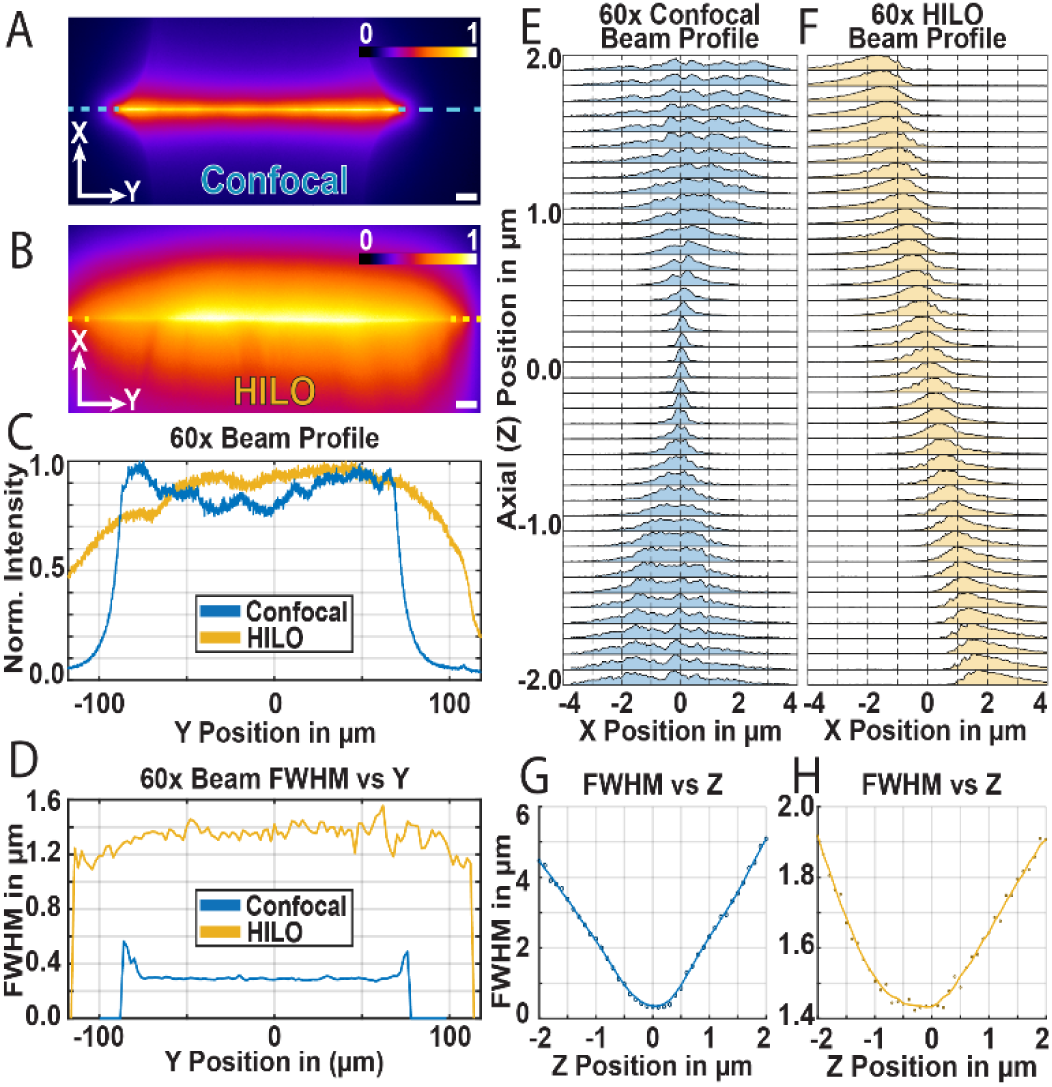
Characterization of the beam profiles for the HIST and line-scanning confocal excitation beam generated from the custom module with the 60X objective. **(A)** Line-scanning confocal excitation line profile from a fluorescent slide (Thorlabs – FSK4). **(B)** HILO line profile from a fluorescent slide (Thorlabs – FSK4). The scale bar for (A) and (B) is 10 µm. **(C)** Excitation beams’ intensity profile across the Y axis from (A) and (B). **(D)** The measured beam’s FWHM along the Y axis from a fluorescent bead sample moved across the excitation line. **(E)** Line-scanning confocal beam profiles along the Z (axial) direction. **(F)** HILO beam profile along the Z (axial) direction. **(G)** Line-scanning confocal beam FWHM along the Z (axial) direction. **(H)** HILO beam FWHM along the Z (axial) direction.

### 2.2 Experimental characterization of the excitation profiles

For each mode and magnification, we used uniformly fluorescent slides to visualize the beam cross section (**Figure 2A-C**) and diffraction-limited beads to probe the beam profile across a grid of points in 3D space (**Figure 2D-H**). Below, we provide characterization under 60x magnification with comparable plots for the 40x and 100x objectives provided as supplementary figures (**Supplementary Figure 2, 3**). The line-scanning confocal profile had a diffraction-limited focus (**Figure 2A, D**) of roughly 0.35 µm, closely matching the expected 0.36 µm design target. The beam maintained a tight focus and uniform intensity profile across a ∼150 µm region (**Figure 2C, D**) that corresponds to the effective field-of-view that can be used along the Y-dimension, while showing sharp increases in FWHM and intensity profile at the edges (**Figure 2C, D**). This follows the expected profile from a large-angle Powell lens and, as noted above, is substantially smaller than the expected 250 µm design width (**Supplementary Table 2**). This is due to challenges in effectively collimating the beam after the Powell lens, which leads to non-uniform illumination profiles when placed at the original design location. As a result, we chose to balance illumination uniformity and field of view by moving the Powell lens closer to the relay optics, effectively clipping the most divergent rays of the fan. On the other hand, the HILO beam profile, generated with a smaller-angle Powell lens and a different set of relay optics, showed a less confined beam with the expected FWHM of ∼1.4 µm (**Figure 2D**) and closely matched the targeted 250 µm Y field-of-view at about 227.5 µm as the measured effective field-of-view (**Figure 2C**).

In the axial dimension, the line-scanning confocal beam showed a well-confined profile with a narrow waist centered at the focus with a Rayleigh length of ∼ 0.5 µm, and symmetric divergence above and below the focus, reflecting the strong optical confinement expected for confocal excitation (**Figure 2E, G**). By comparison, the HILO beam produced a larger FWHM at the focus with a longer Rayleigh length of ∼3.5 µm and a beam angle of 34.13° relative to the detection plane, which is consistent with prior HIST implementations using similar systems (**Figure 2F, H**)^11,12^.

### 2.3 Experimental characterization of the point spread and optical transfer functions

After characterizing the illumination profiles, we next characterized the overall imaging performance using the PSF and OTF under varying rolling shutter integration widths. The excitation modes introduced in the custom modules show a clear improvement in terms of optical sectioning compared to widefield (**Figure 3A-D (i)**), with progressively improved sectioning with narrower integration widths approaching 1 Airy unit, consistent with its function as a virtual confocal slit. The improvement in optical sectioning is also apparent in the calculated OTFs, where both HIST and line-scanning confocal modes demonstrated expanded frequency support in the axial direction and improved lateral frequency support along the k_x_ axis (parallel to the line scanning direction) (**Figure 3A-D (ii-iv)**).

**Figure 3.**
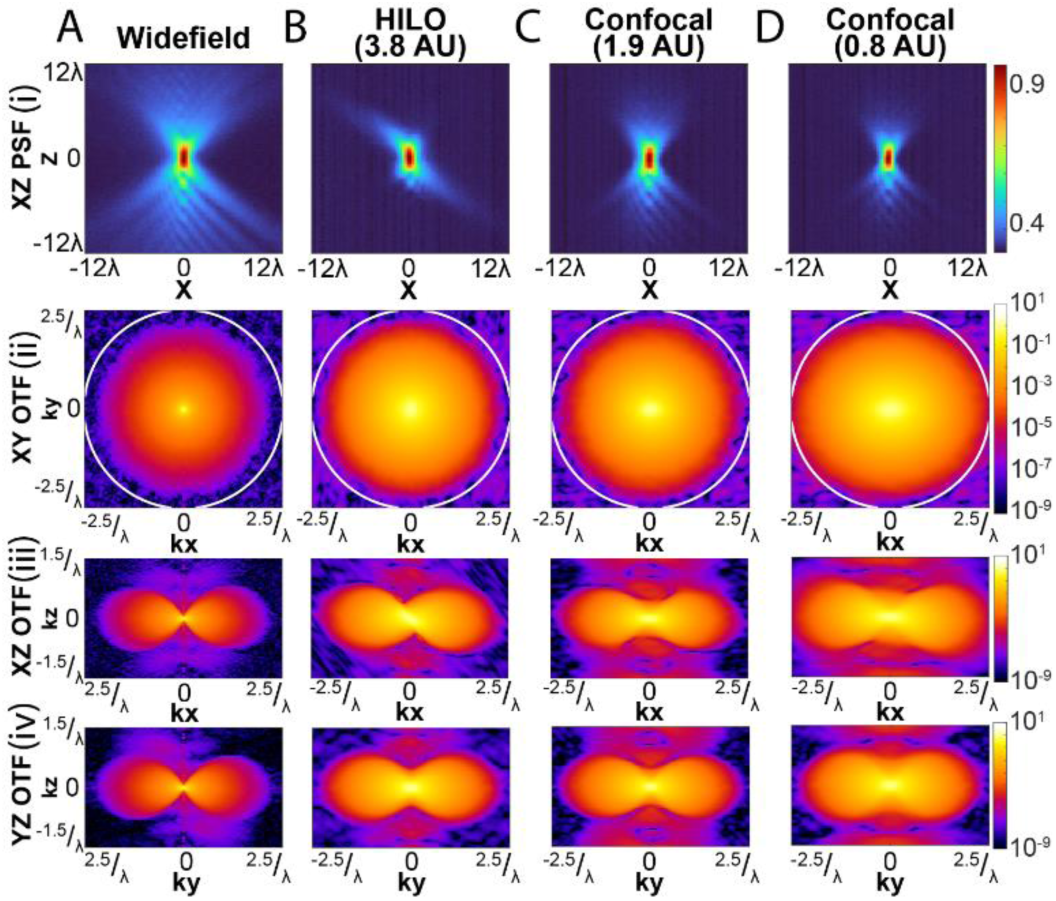
PSF and OTF comparison of the custom module’s different excitation methods with the 60X objective. **(A)(i)** Widefield excitation scheme characterized by its experimentally measured XZ PSF. **(ii)** XY OTF slice computed from the experimentally measured PSF. **(iii)** XZ OTF slice computed from the experimentally measured PSF. **(iv)** YZ OTF slice computed from the experimentally measured PSF**. (B-D)** are the same as (A) for HIST with a rolling shutter/slit width of 3.8 Airy units and line-scanning confocal with rolling shutter/slit widths of 1.9 and 0.8 AU.

The ability to tune the rolling shutter integration width “on-the-fly” with no mechanical components allows the user to balance optical sectioning and signal collection efficiency, in addition to being able to tune this parameter for different magnifications. Detailed quantifications for the 40x and 100x objectives are available in (**Supplementary Figures 2-5**). Finally, while the system can switch between HIST and line-scan confocal without realignment or replacement of any optical components, we also designed a third module that can convert the system to TIRF illumination by removing the Powell lens and exchanging the relay lenses in the HIST beam shaping module (**Supplementary Figure 6**). In this configuration, the system demonstrated uniform TIRF illumination across a ∼120 µm field of view while confining the illumination to the evanescent field above the coverslip, further increasing the range of experiments that can be performed with the system.

### 2.4 Experimental demonstration

We next demonstrated the capabilities of the system’s two excitation modes on representative biological samples. At 60x magnification, the line-scanning confocal mode provided improved optical sectioning compared to widefield on multi-color immunofluorescence datasets taken of CAR-T and CD19-expressing target cells (**Figure 4A, B**). In live cells, the line-scanning confocal mode can also be utilized for fluorescence recovery after photobleaching (FRAP) experiments to measure protein mobility within 3D specimens (**Supplementary Figure 7**). For single-molecule and super-resolution imaging, HIST illumination provides a large field and low background, which we demonstrate by acquiring two-color DNA-PAINT datasets of Lamin AC and H3K27Ac (**Figure 4C-E**). These results demonstrate the capability of the custom module to support a wide range of fluorescence imaging experiments, spanning multicolor volumetric imaging, live-cell measurements of protein dynamics, and super-resolution microscopy within a single platform.

**Figure 4.**
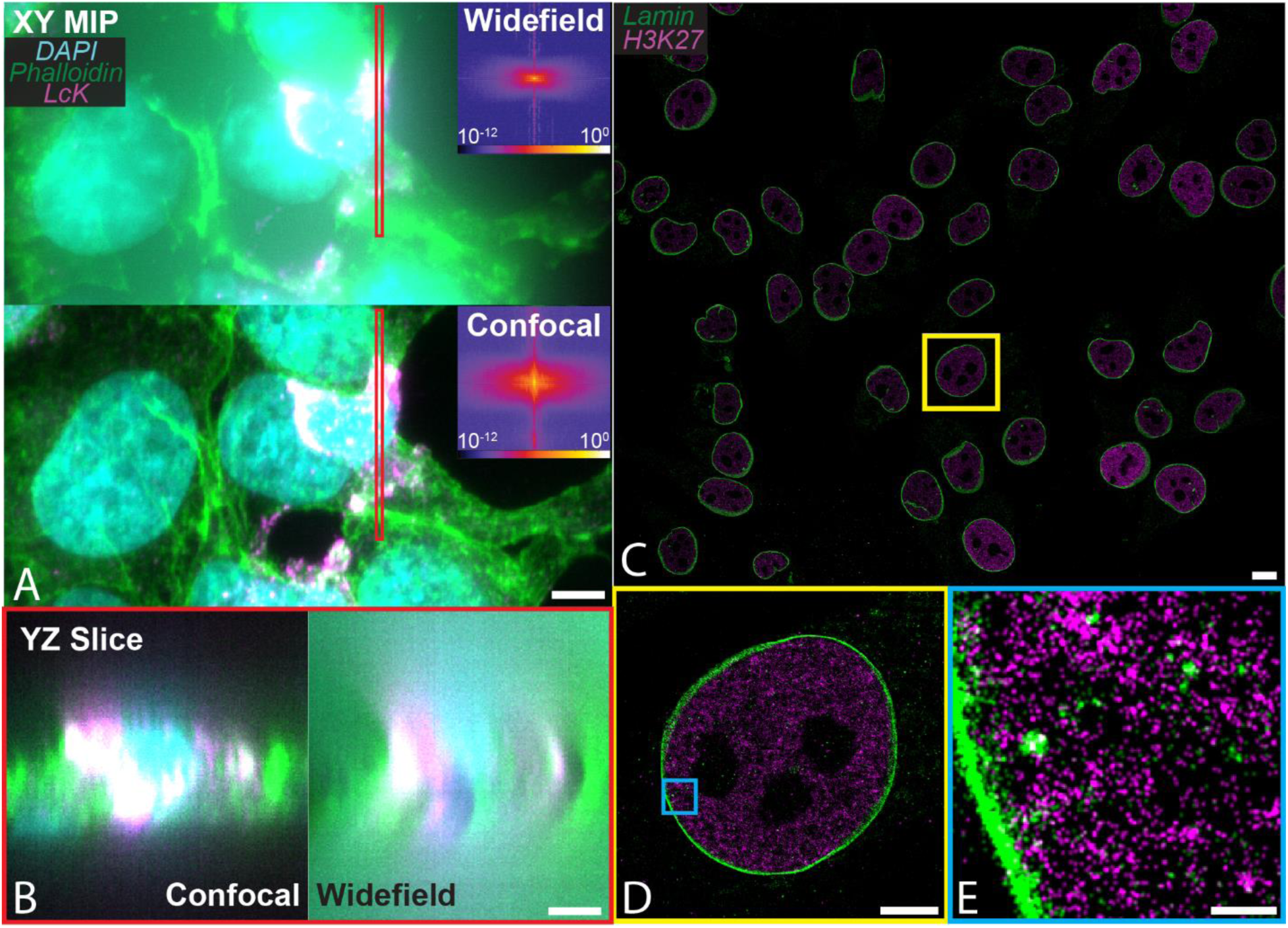
Fixed cell representative images obtained using the custom excitation module with the 60x objective. **(A)** Representative XY MIP images of immune synapses formed between a CAR-T and CD19 expressing target cells showing Lck (magenta), nuclei (cyan), and actin (green). The insets are the corresponding Fourier transform of the actin channel in XZ. **(B)** YZ slice of the highlighted red region from (A). The scale bar is 5 µm. **(C-E)** DNA-PAINT representative images of Lamin (green) and H3K27Ac (magenta) in SKMEL5 melanoma cells. The scale bar is 20 µm for (D), 10 µm for (E), and 100 nm for (F).

The ability to alternate between line-scanning confocal and HIST imaging within a single instrument also enabled the integration of automatic mode switching for more complex experiments. As an example, we alternated between confocal imaging and single-particle tracking to quantify single-molecule dynamics and the spatial organization of chromatin in the nucleus (**Figure 5**). Line-scanning confocal imaging provided optically sectioned three-dimensional information of the nuclear architecture (**Figure 5A, B**), allowing chromatin organization to be resolved and segmented into distinct chromatin classes (**Figure 5B**), while HIST imaging enabled high-contrast single-particle tracking (SPT) measurements of nucleoplasmic proteins (**Figure 5A**).

**Figure 5.**
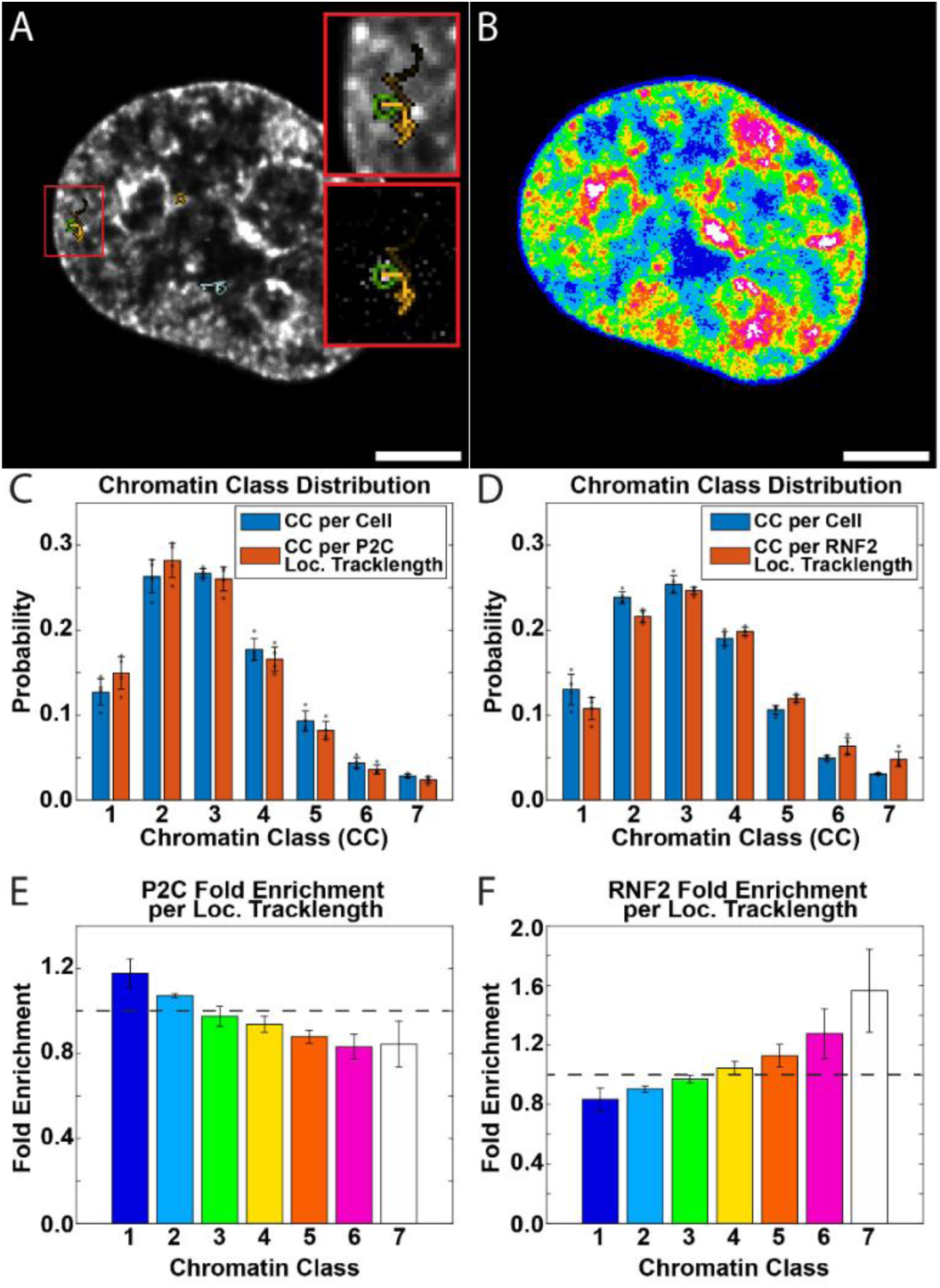
Correlative confocal imaging and HIST-assisted single-particle tracking (SPT) reveal preferential localization of P2C and RNF2 trajectories across chromatin classes. **(A)** Representative confocal image of a nucleus with overlaid SPT trajectory (yellow track) and magnified view of the corresponding track (top: zoom in of the overlaid track, bottom: zoom in of the raw track data). **(B)** Visualization of the chromatin class segmentation into 7 different classes, ranging from low- (1) to high-density (7) class. **(C, D)** shows the chromatin class distribution on P2C SPT data and their relative enrichment for each chromatin class. **(E, F)** shows similar data for RNF2. Scale bars are 5 µm for (A, B).

Alternating between both imaging modes in one experiment, with a switching time of ∼2 seconds, allowed us to quantify how the localization and molecular dynamics changed as a function of chromatin density. For example, we found that fluorescently tagged RNA polymerase II molecules were more likely to be found in lower chromatin density classes (**Figure 5C, E**), whereas ring finger protein 2 (RNF2), a component of the polycomb repressive complex one (PRC1), was enriched in denser chromatin compartments (**Figure 5D, F**). This multimodal workflow demonstrates the expanded experimental capability and convenience of an integrated custom module, providing optically sectioned volumetric context alongside dynamic single-molecule measurements. Altogether, these results show the capabilities of the integrated custom module as a versatile platform for biological discovery while removing the need to transfer between instruments to use different modalities.

## 3 Discussion

While line scan confocal and HIST are established techniques with clear advantages for different imaging experiments^10,14^, to our knowledge, they have not yet been integrated into a single platform. Compared to prior implementations of HIST imaging^10–12^, the addition of line-scanning confocal transforms the system from one that is optimized primarily for single-molecule imaging to one that can be applied to a broad spectrum of biological questions. Here, we demonstrate how this can be accomplished with the addition of only a few optical components and the necessary hardware to toggle between different beam-shaping modules. By sharing much of the hardware and optical path, adding this new functionality increases the cost by only 13% compared to a comparably equipped HIST-only system (laser assembly and custom hardware, not including the Nikon scope body and peripherals)^12^. Through careful design considerations, we also demonstrate how the same system can be implemented across several different magnifications, further increasing its versatility for biological imaging.

In the confocal mode, the sCMOS rolling shutter functions as a virtual confocal slit which can be easily adjusted programmatically with no mechanical parts. However, there are trade-offs to this approach. Compared to a conventional descanned confocal system with shared scanning and detection paths, the sCMOS-based approach requires precise alignment and synchronization between the scanning optics and camera readout. For example, in our highest resolution imaging condition (100x magnification, 1.35 NA, 0.5 Airy unit integration width), the projected confocal slit is only 260 nm wide at the sample. In our experience, to achieve uniform illumination, the excitation line must stay registered with this stripe to within roughly 50 nm while sweeping across the entire 150 µm FOV. This can become challenging at high scan speeds due to the inertia of the galvo and physical vibrations from the camera fan. In our system, we accounted for this by choosing a small (3 mm) galvo with low inertia and water-cooling the camera, but future implementations using larger galvos or non-water-cooled cameras would need to take this into account. However, we note that the precision of the alignment and timing are substantially relaxed for lower magnifications and larger rolling shutter integration widths, making this the most extreme example.

Overall, we envision that this work will be a useful contribution for research labs that are considering HIST illumination for single-molecule imaging and wish to add additional functionality to their instruments. It may also increase the possibility that such systems can be integrated into imaging cores where a single instrument must cater to a diverse range of research labs and applications. In the future, the ability to automatically toggle between different imaging modes could also contribute to the growing field of “smart microscopy” wherein the microscope is programmed to react to the images acquired from the biological specimen^15^.

## 4 Materials and Methods

### 4.1 Modified Nikon microscope setup

We designed the HIST/confocal module to integrate with a Nikon Eclipse Ti2-E microscope body equipped with a motorized stage, piezo-driven z-drive (Queensgate QGNPS083-NI) for precise and fast z-stack acquisition, and an Okolab live cell chamber (Okolab H301) to perform extended live cell imaging. The custom illumination module mounts onto the back port of the microscope through a motorized Nikon Ti2-LAPP module that allows switching between the custom module and an existing epifluorescence LED light source (Nikon D-LEDI). After entering the microscope, the excitation beam from the custom module is reflected off a quad-band dichroic mirror (Chroma ZT405/488/561/640rpcv2-UF2) and directed onto the pupil of one of three objective lenses (Nikon MRD73400, MRD73950, or MRY10060). The sample fluorescence emission is collected by the objective, passes back through the dichroic, is filtered by a quad-notch emission filter (Semrock NF03-405/488/561/635E-25), and is imaged onto an sCMOS camera with rolling shutter acquisition and “light sheet readout mode” (Hamamatsu C14440-20UP)^16^.

### 4.2 Line-scanning confocal/HIST custom module

The custom illumination module utilizes a previously described laser combiner^12,17^ where 4 laser lines (TOPTICA IBEAM-SMART-405-S-BZ, MPB 2RU-VFL-P-500-488-B1R, MPB 2RU-VFL-P-2000-560-B1R, MPB 2RU-VFL-P-2000-642-B1R) are made colinear through a series of dichroic mirrors before passing through an acousto-optic tunable filter (AOTF - AA Optoelectronic, AOTFnC-400.650-CPCh-TN) for wavelength selection and power modulation. The AOTF output is coupled into an optical fiber (kineMATIX-P2 and kineFLEX-HPV) and transmitted to the custom excitation module. The custom module is a combination of bespoke parts and off-the-shelf components to ensure precision, a compact form factor, and a convenient assembly process. It consists of a custom-machined base plate to mount various optomechanical components such as the optical fiber coupler, toggle mirrors (Thorlabs MRA-P01 right angle silver mirrors on SmarAct SLC stages), Powell lenses (Laserline Optics LOCP-8.9R10-0.8 and LOCP-8.9R30-1.8), and relay lenses. The scanning galvo mirror (Thorlabs GVS201), scan lens (Thorlabs CLS-SL), and tube lens (Thorlabs TTL200MP) are located on a platform extending perpendicular from the main base plate to accommodate the design geometry while keeping a small footprint. The complete optical path, CAD model, and parts lists are outlined in Figure 1 and in more detail within the supplementary materials. The custom assembly, camera, and Nikon microscope stand are all controlled via an FPGA (NI, USB-7845R OEM) and a modified version of the MOSAIC microscope software written in LabVIEW 2022 Q3 (64-bit) previously published in^17^.

### 4.3 Confocal excitation beam and rolling shutter alignment

The confocal excitation beam needs to be precisely aligned with the rolling shutter (proxy for a confocal slit) of the camera to obtain optimal performance. To accomplish this, we removed the emission filter and imaged the back reflection of the excitation beam from the coverslip-water/oil interface onto the camera. We lowered the laser power and used a narrow (∼4 pixels) rolling shutter width to minimize potential saturation of the camera from the back-reflected beam. We then adjusted the galvo offset voltage to center the back-reflected beam on the camera chip.

The galvo scanning and rolling shutter speed were matched by optimizing the galvo’s volts-to-microns conversion factor to match the exact magnification of the system for each objective. The exact magnification of each objective on the system was obtained by moving a fluorescent bead in set amounts using the Nikon XY stage and performing a linear fit for the slope of the microns/pixel measurements. Timing pulses from the camera (one pulse per row readout) in HSYNC mode were used to synchronize the galvo sweep timing.

### 4.4 Optical characterization

To qualitatively observe the beam profile for the line-scanning confocal and HIST modes, we imaged a homogeneous fluorescent slide (Thorlabs – FSK4) illuminated using the 560 nm laser. As this approach convolves the excitation pattern with the emission PSF, we also utilized sub-diffraction 100 nm fluorescent bead samples (Invitrogen FluoSpheres F8801) to more directly measure the excitation profiles. We measured the width of the confocal and HIST beams by scanning the excitation beam along the X-axis in 50 nm steps across a static bead and integrating the camera counts from a small 3 x 3-pixel region centered on the bead at each galvo location. In this case, the fluorescent bead acts as a sub-diffraction reporter whose fluorescence is proportional to the local excitation intensity. We then fit the plots of the bead intensity vs. beam position to a Gaussian function to determine the beam FWHM. We characterized the uniformity of this measurement across the Powell lens extension (Y-axis, long axis of the beam) by translating the bead in 2 µm steps in Y. The excitation beam profiles in the axial direction were visualized similarly using a fluorescent bead that was translated in 100 nm steps along the Z-axis using the sample piezo Z stage.

We computed the 3D PSFs by collecting z-stacks of isolated 100 nm fluorescent beads (Invitrogen FluoSpheres F8801) at 50 nm steps. To increase the signal-to-noise ratio, we averaged over 5 images per z-plane and background-subtracted the measured dark current data, obtained by averaging 300 dark current images under the same exposure time. We then re-centered and normalized the PSF intensity before Fourier transforming to acquire the corresponding experimental OTFs for each excitation mode. To aid visual interpretation, we gamma-adjusted and clipped the XZ PSFs for each mode using a consistent gamma value of 0.2 and scaled the display to saturate the top and bottom 1% of the intensity. The OTFs were further processed by calculating the log scale of their absolute value (normalized from their imaginary values) for display purposes.

### 4.5 Cell culture and sample preparation

#### 4.5.1 SK-MEL5 and CAR T cells

The hCD19^+^ SK-MEL5 cell line used in Figure 4A, B was generated by engineering the HLA- A2*^+^* SK-MEL5 melanoma cell line to constitutively express human CD19 and eGFP-FFluc via retroviral transduction. Cells were cultured in RPMI 1640 with L-glutamine (Gibco 11875-093) supplemented with 10% fetal bovine serum (FBS; Gemini 512450) at 37°C, 5% CO*_2_*, and 100% humidity. SK-MEL5 cells were plated on fibronectin-coated 8-mm glass coverslips in individual wells of a 6-well plate and cultured to 70% confluency. Viral vectors for T-cell transduction were transiently produced by transfecting HEK 293T cells (ATCC) with GeneJuice according to the manufacturer’s protocol (Sigma). Retroviral constructs included the RD114 envelope plasmid, MoMLV gag-pol (PegPam3-e plasmid), and a CD19 CAR vector (FMC63 clone with G4S linker). Lentiviral constructs included the VSVG envelope plasmid (pMD2.G; Addgene), second-generation lentiviral packaging plasmid psPAX2 (Addgene), and an in-house-designed CD19 CAR vector containing a FLAG tag for detection (VectorBuilder). Virus-containing supernatants were sterile-filtered through 0.45-μm syringe filters, aliquoted, snap-frozen in liquid nitrogen, and stored at −80°C until use. CAR T cells used in Fig. 4a were generated as previously described^18,19^. Briefly, frozen human PBMCs isolated from buffy coats were activated on anti-CD3/anti-CD28-coated plates for 2 days, followed by stimulation with recombinant human IL-2 (PeproTech 200-02) for 1 day. On the day of transduction, the virus was thawed and concentrated 20-fold by centrifugation in Amicon filter units (100-kDa cutoff). Concentrated virus and 2 × 10*^6^* activated PBMCs (50 μL total volume) were seeded onto 24-well alginate scaffolds (Lenti-X sponges, Takara Bio) for transduction. Following transduction, cells were cultured in complete medium with IL-2 for 72 h, then isolated from the sponge, validated for CAR expression by flow cytometry, aliquoted, frozen, and stored for future use. CAR T cells were cultured in a 50:50 mixture of Click’s medium (HyClone 502892160) and RPMI 1640 containing 1% GlutaMAX (Gibco 35050061), supplemented with 10% FBS, at 37°C, 5% CO*_2_*, and 100% humidity. Cells were passaged and stimulated with 250 IU IL-2 every 48 h and maintained at 0.5 × 10^6 cells/mL for a maximum of 3 weeks to prevent exhaustion.CD19 CAR T cells were added to SK-MEL5-coated coverslips at a 1:1 effector-to-target ratio and incubated for 2 h at 37°C. Co-cultures were fixed with 4% PFA in 1× PBS for 20 min, followed by three 5-min washes in 1× PBS. Cells were permeabilized with 0.2% Triton X-100 in 1× PBS for 15 min at room temperature (RT), washed with 1× PBS, and blocked overnight at 4°C with 10% normal goat serum (Thermo). Samples were then incubated with rabbit anti-LCK primary antibody (Cell Signaling #2984; 1:1000) for 2 h at RT, washed three times in 1× PBS for 5 min each, and incubated with Alexa Fluor 647 secondary antibody (Thermo A-21245; 1:500) for 2 h at RT. Following three additional 5-min washes in 1× PBS, samples were labeled with Phalloidin 555 (Thermo A-34055; 1:500) and DAPI (Sigma D9542; 1:1000) for 1 h at RT. Coverslips were washed three times for 5 min in 1× PBS, removed from the wells, inverted onto glass slides, and mounted with ProLong Diamond (Thermo P-36961) overnight at RT before imaging.

#### 4.5.2 RNF2 and P2C HaloTag HEK293T cells

RNF2 and P2C HaloTag-tagged cell lines were generated using a previously described method with minor modifications^20^. Briefly, HEK293T cells were transfected with a donor DNA molecule containing the HaloTag marker we wished to knock-in along with a tandem in-frame ribosomal skipping peptide PT2A followed by 3 copies of the nourseothricin resistance marker. At the same time the donor molecule is transfected, a Cas9-containing plasmid is provided along with a plasmid containing a gRNA targeting the C-terminus of our genes of interest and a second gRNA against our donor plasmid. Following transfection, cells were maintained for 7 days in non-selective conditions to allow for integration of the donor construct and sufficient time for the drug selection marker to be expressed. To enrich for cells containing the correctly integrated HaloTag cassette, nourseothricin was subsequently applied for selection at 100 mg/ml. Resistant cells were expanded to establish stable RNF2- and P2C-HaloTag-tagged cell lines. Correct integration of the HaloTag was verified by PCR, confirming the in-frame fusion of the HaloTag with our genes of interest. The expression and localization of the HaloTag-fused proteins were further validated by immunofluorescence.

The HEK293T-Halo-P2C and HEK293T-Halo-RNF2 were maintained in DMEM (Thermo Fisher Scientific, 11965118) and supplemented with 10% fetal bovine serum (GeminiBio, S12450) and 10% (v/v) penicillin-streptomycin (Thermo Fisher Scientific, 15140122). Cells were regularly screened for mycoplasma contamination using imaging and isothermal PCR techniques. For live imaging, cells were plated in 24-well glass-bottom plates with high-performance #1.5 cover glass (Cellvis P24-1.5H-N).

HaloTag ligands (HTL) were dissolved in anhydrous DMSO (Thermo Fisher Scientific, D12345) to reach a final concentration of 1 mM. Stock solutions were then aliquoted and stored at −20 °C; a freshly thawed aliquot was used immediately for each experiment. The ligand stock solution was diluted into complete growth medium, and cells were incubated for 1 h. To achieve an appropriate labeling density for single-particle tracking, HTL stock solution was diluted to a final working concentration of 0.1-0.15 nM. To achieve saturating labeling of HTL for FRAP, the stock solution was diluted to a final working concentration of 100 nM.

One day before imaging, 24-well glass-bottom plates were incubated with 10ug/mL fibronectin (Sigma-Aldrich, F2006) for 20 min and washed with PBS (Thermo Fisher Scientific, 10010049). Cells were then plated at a density of ∼ 4.2 × 10^4 cells/cm². On the day of imaging, the cells were incubated in growth medium containing either 0.1-0.15 nM (SPT) or 100 nM (FRAP) of the HaloTag-Ligand JaneliaFluor 646 and a 500X dilution of the manufacturer-recommended stock concentration of SPY505 (Cytoskeleton, CY-SC101) for 1 h. Following incubation, growth medium was replaced with dye-free Fluorobrite DMEM medium (Thermo Fisher Scientific, A1896701) supplemented with 10% (v/v) penicillin-streptomycin (Thermo Fisher Scientific, 15140122)4.6 Imaging protocol and analysis pipeline

Detailed imaging settings for each dataset are described in **Supplementary Table 3**. For the multi-color 3D fixed cell images, we compared line-scanning confocal and widefield excitation by sequentially acquiring each color channel at each z-plane. We obtained the Fourier spectrum from these images by isolating the phalloidin channel and performing FFT on the corresponding 3D image stack for each excitation mode. To minimize edge artifacts, we first processed the stacks using the *perdecomp_3d* function from the DIPImageToolbox^21^ and then mirror- and zero-padded the data before the Fourier transform. Both confocal and widefield spectra were processed identically.

FRAP datasets were acquired by utilizing the line-scanning confocal mode of the custom module. The experiment starts with imaging the sample at 25 ms exposure time for 13 s (325 ms cycle time, 40 images), and is followed by a bleaching step for 1 s at 96.7 mW (642 nm laser, measured at the objective collar) by dithering the confocal line beam in a small range (∼1 µm) at the center of the FOV, before imaging the fluorescence recovery for 325 seconds every 325 ms. The analysis is done by first segmenting the nuclei manually using Amira3D 2023 to generate the nuclei masks. The width of the bleached stripe was defined based on the set experimental settings and adjusted for each nucleus over time based on the movement of the nucleus’s center of mass obtained from the generated nuclei masks. The FRAP recovery curves were generated by measuring the fluorescence intensity of the bleached area for each affected nuclear mask, subtracting the immediate post-bleach intensity in the bleached area, and normalizing by the pre-bleach intensity. The resulting FRAP curves were fit to a two-component exponential recovery curve, which was used to estimate the half-recovery time.

For the DNA-PAINT experiment, we sequentially acquired each channel. Localizations were identified from raw images using the SMAP package in MATLAB^22^ and processed as described in^12^.

For the combined confocal and SPT switching experiment, we deconvolved the confocal images using a Richardson-Lucy-based GPU-accelerated deconvolution algorithm^23^ with an OTF calculated from an average of 3 experimental PSFs after experimental dark current noise subtraction. The deconvolved image stacks were used together with Cellpose-SAM^24^ to generate nuclear masks. These masks were then used to select the raw pixel values that were used to classify chromatin classes. Chromatin classification was performed by normalizing the pixels from each cell to clip the top and bottom 1% and then binning the intensity values into 7 different histogram bins/classes distributed between zero and one. The generated classes were then used to classify the SPT localizations obtained with the HIST excitation mode. We used TrackMate^25^ to localize and track individual molecules from the HIST images using the following settings: for object detection, the Difference of Gaussians (DoG) detector was used, with an estimated object diameter of 0.5 μm, a quality threshold of 10.5, and sub-pixel localization. For tracking, the simple Linear Assignment Problem (LAP) tracker was used with a linking max distance of 0.7 µm, a gap-closing max distance of 1.5 μm, and a gap-closing max frame gap of 5. We then sorted each localization into a chromatin class by comparing its location to the corresponding chromatin classification image obtained immediately before the HIST acquisition. We then compared the enrichment of molecules in each class to what would be expected from a randomly distributed localization within the nucleus.

## Code, Data, and Materials

All data supporting this paper are available from the corresponding authors upon request. The LabVIEW software implementation to control the microscope is available through a research license agreement with the Howard Hughes Medical Institute.

## Acknowledgments

This work was supported by NIH grant 1R35GM158040 to W.R.L. W.R.L. acknowledges additional support from the Packard Fellowship for Science and Engineering. We thank Yevgeny Brudno and Gianpietro Dotti for sharing the CAR-T and CD19+ MKMEL5 melanoma cells, respectively.

