## Supplementary Material for "A custom two-in-one HIST and line-scanning confocal excitation module"

*Supplementary Figure List*

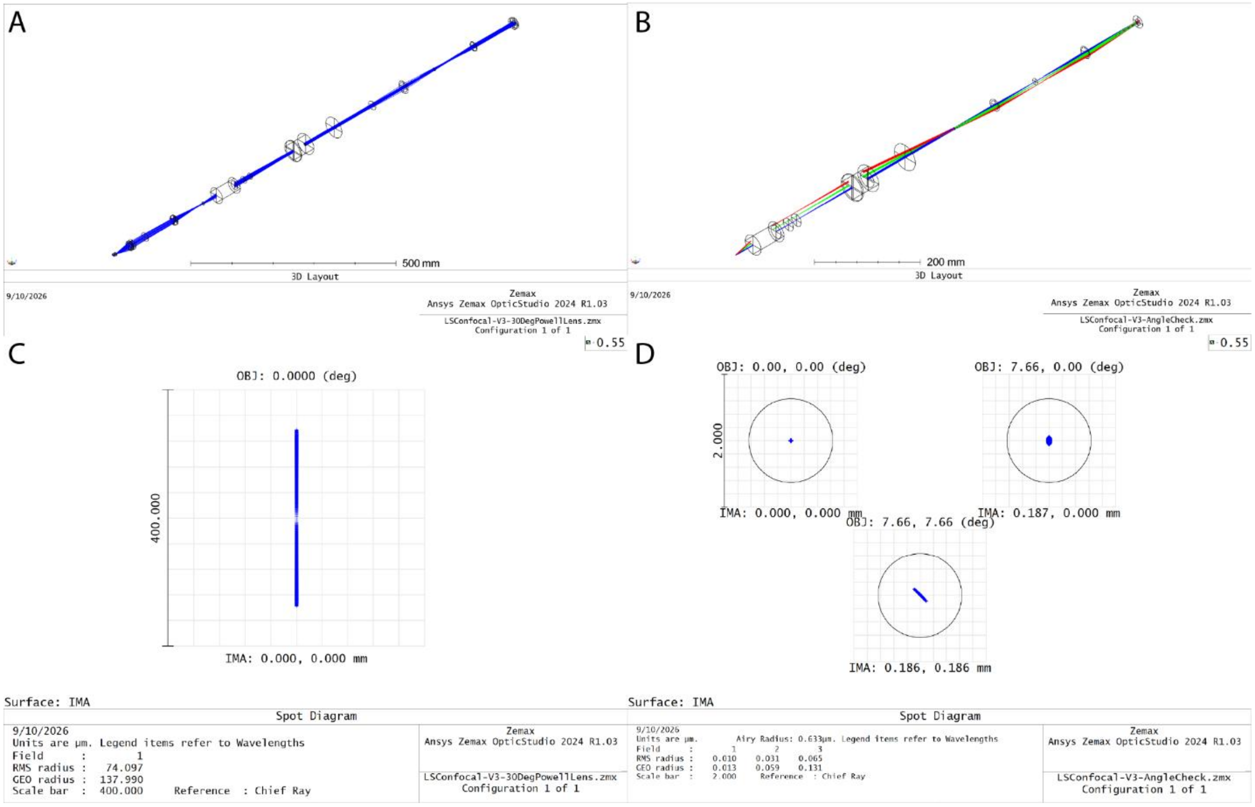

**Supplementary Figure 1.** Zemax simulation of the custom line-scanning confocal and HIST module. (A) 3D layout of the entire optical path in the custom module (from the Powell lens to the sample). (B) 3D layout of the post-galvo scanning optical path of the custom module, with additional rays corresponding to the edges of the targeted FOV (i.e. 375 µm for a 40x objective). (C) Spot diagram of the generated beam on the sample plane (assuming centered beam throughout the system). (D) Spot diagrams of a focused beam (assuming collimated light post-galvo mirror) on the sample plane at the center and edges of the targeted FOV. Nikon objective and additional lenses are simulated as perfect lenses in these tests.

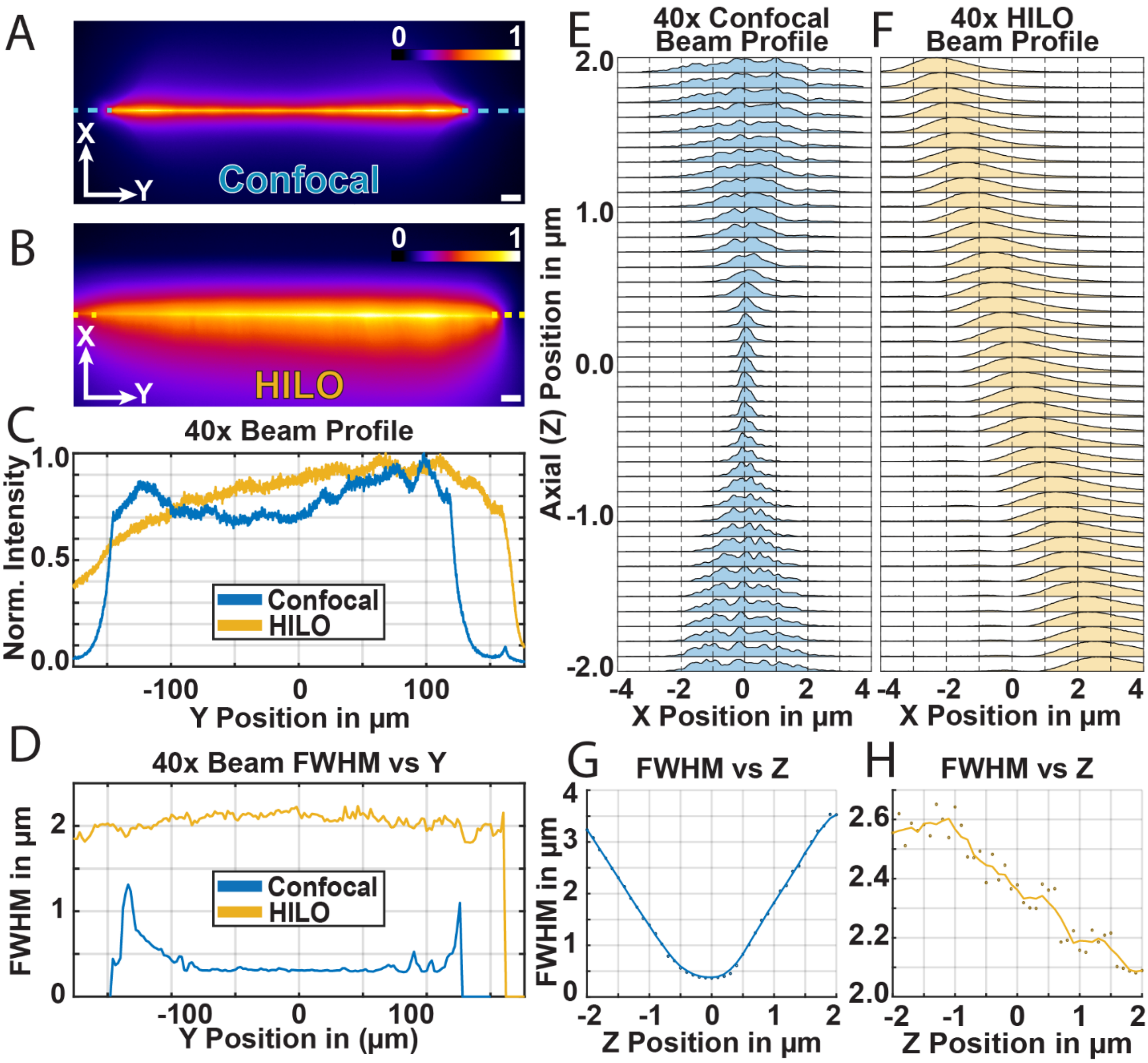

**Supplementary Figure 2.** Characterization of the beam profiles for the HIST and line-scanning confocal excitation beam generated from the custom module with the 40x objective. **(A)** Line-scanning confocal excitation line profile from a fluorescent slide (Thorlabs – FSK4). **(B)** HILO line profile from a fluorescent slide (Thorlabs – FSK4). The scale bar for (A) and (B) is 10 µm. **(C)** Excitation beams' intensity profile across the Y axis from (A) and (B). **(D)** The measured beam's FWHM along the Y axis from a fluorescent bead sample moved across the excitation line. **(E)** Line-scanning confocal beam profiles along the Z (axial) direction. **(F)** HILO beam profile along the Z (axial) direction. **(G)** Line-scanning confocal beam FWHM along the Z (axial) direction. **(H)** HILO beam FWHM along the Z (axial) direction.

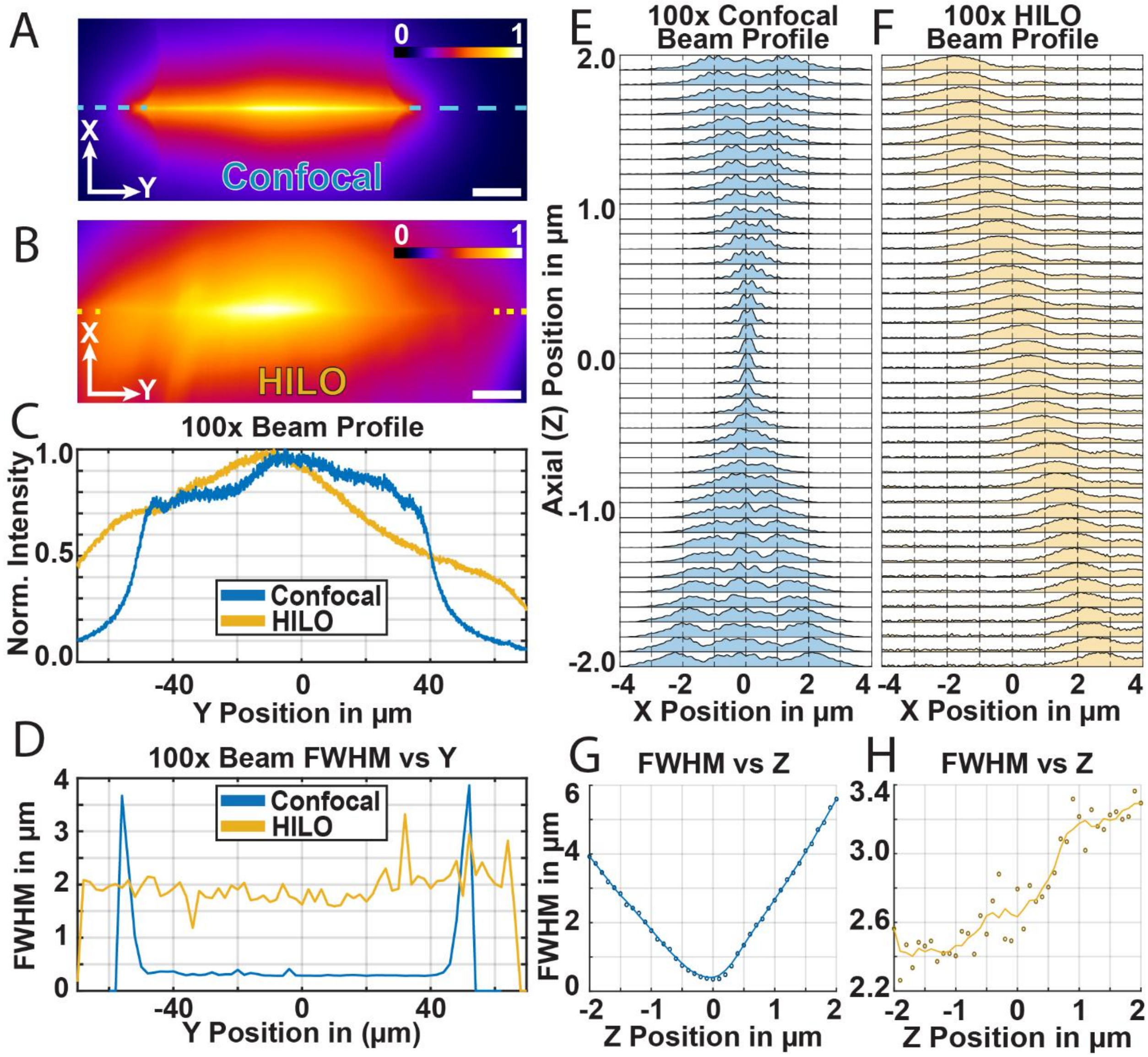

**Supplementary Figure 3.** Characterization of the beam profiles for the HIST and line-scanning confocal excitation beam generated from the custom module with the 100x objective. **(A)** Line-scanning confocal excitation line profile from a fluorescent slide (Thorlabs – FSK4). **(B)** HILO line profile from a fluorescent slide (Thorlabs – FSK4). The scale bar for (A) and (B) is 10 µm. **(C)** Excitation beams' intensity profile across the Y axis from (A) and (B). **(D)** The measured beam's FWHM along the Y axis from a fluorescent bead sample moved across the excitation line. **(E)** Line-scanning confocal beam profiles along the Z (axial) direction. **(F)** HILO beam profile along the Z (axial) direction. **(G)** Line-scanning confocal beam FWHM along the Z (axial) direction. **(H)** HILO beam FWHM along the Z (axial) direction.

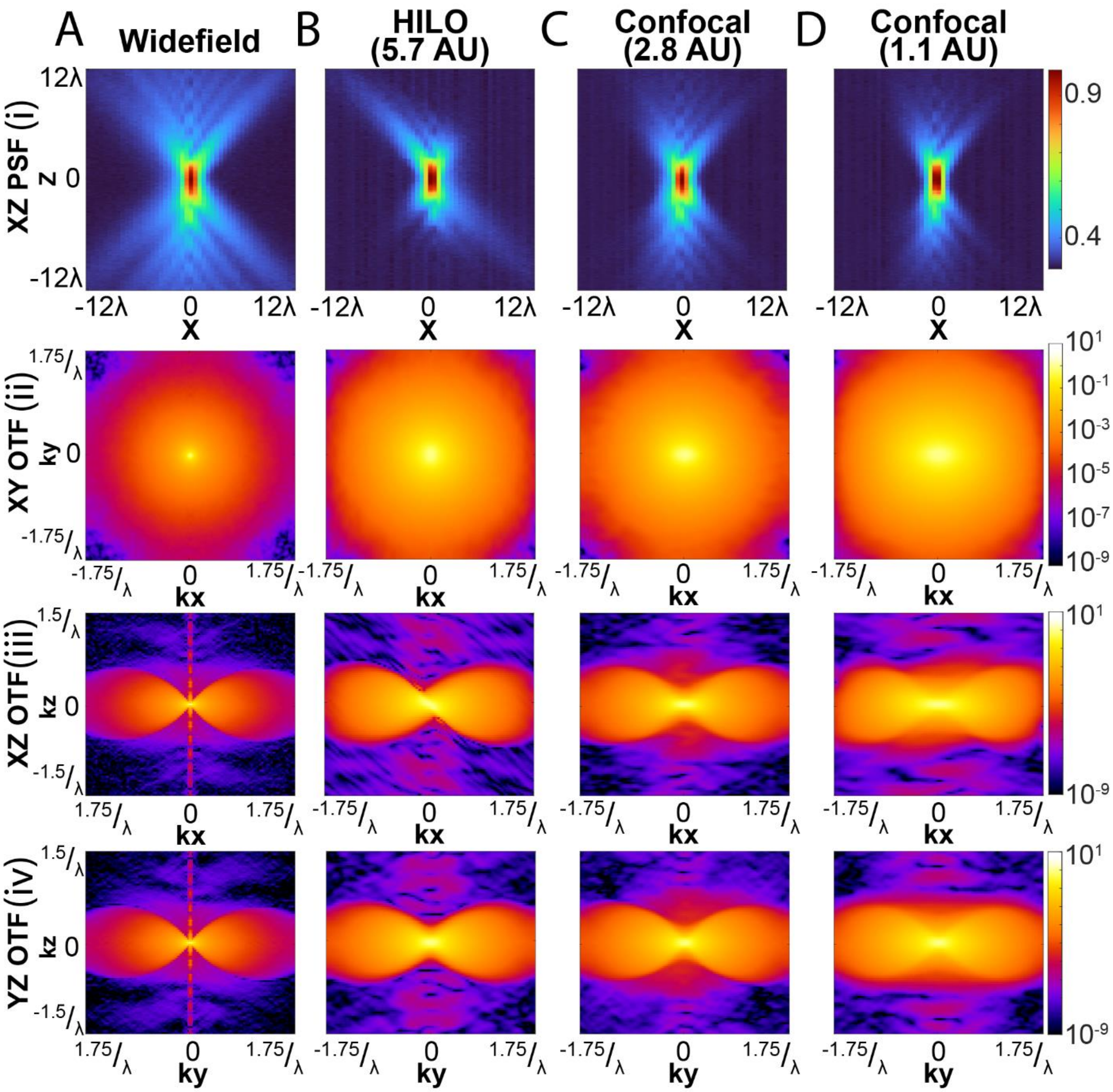

**Supplementary Figure 4.** PSF and OTF comparison of the custom module's different excitation methods with the 40x objective. **(A)(i)** Widefield excitation scheme characterized by its experimentally measured XZ PSF. **(ii)** XY OTF slice computed from the experimentally measured PSF. **(iii)** XZ OTF slice computed from the experimentally measured PSF. **(iv)** YZ OTF slice computed from the experimentally measured PSF. **(B-D)** are the same as (A) for HIST with a rolling shutter/slit width of 5.7 Airy units and line-scanning confocal with rolling shutter/slit widths of 2.8 and 1.1 AU.

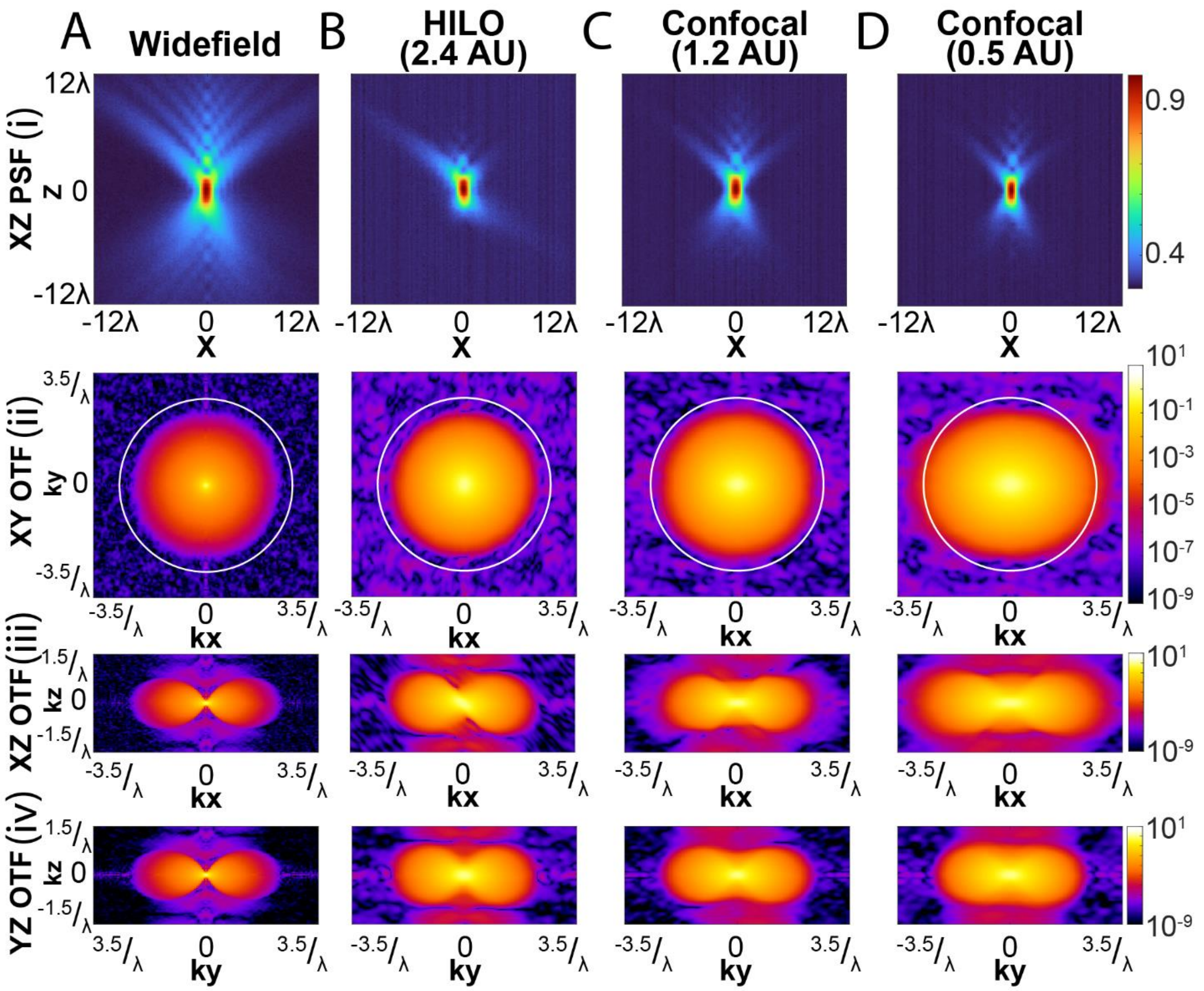

**Supplementary Figure 5.** PSF and OTF comparison of the custom module's different excitation methods with the 100x objective. **(A)(i)** Widefield excitation scheme characterized by its experimentally measured XZ PSF. **(ii)** XY OTF slice computed from the experimentally measured PSF. **(iii)** XZ OTF slice computed from the experimentally measured PSF. **(iv)** YZ OTF slice computed from the experimentally measured PSF. (B-D) are the same as (A) for HIST with a rolling shutter/slit width of 2.4 Airy units and line-scanning confocal with rolling shutter/slit widths of 1.2 and 0.5 AU.

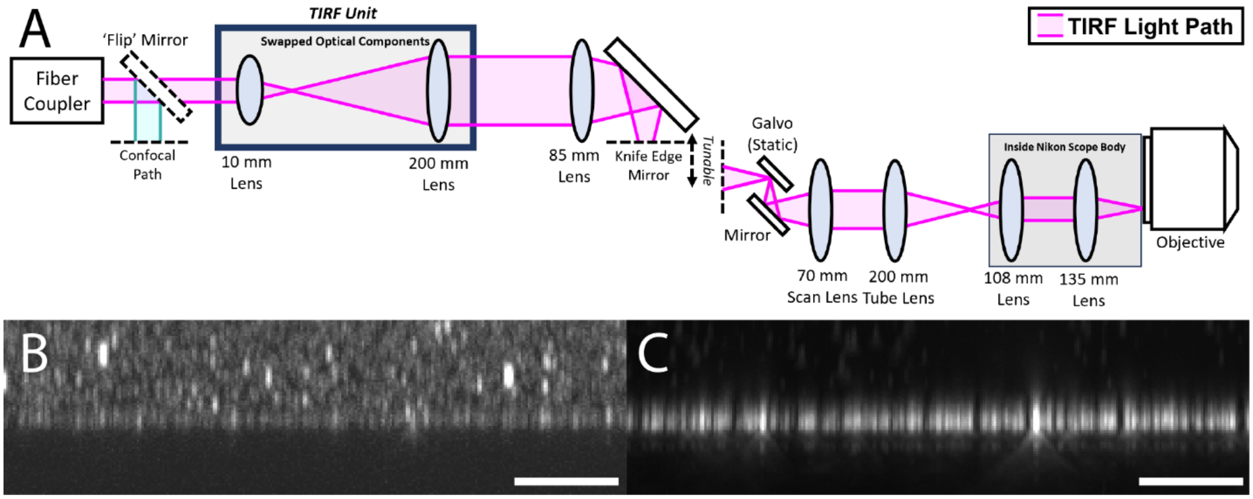

**Supplementary Figure 6.** Insertable TIRF module**. (A)** Optical path of the custom module's HIST path converted into TIRF by swapping the optical elements for the TIRF unit. **(B)** Widefield XZ MIP of fluorescent beads (Invitrogen FluoSpheres F8801) embedded in a PDMS gel. **(C)** TIRF XZ MIP of the same region as (B). Scale bars are 5 µm for (B, C).

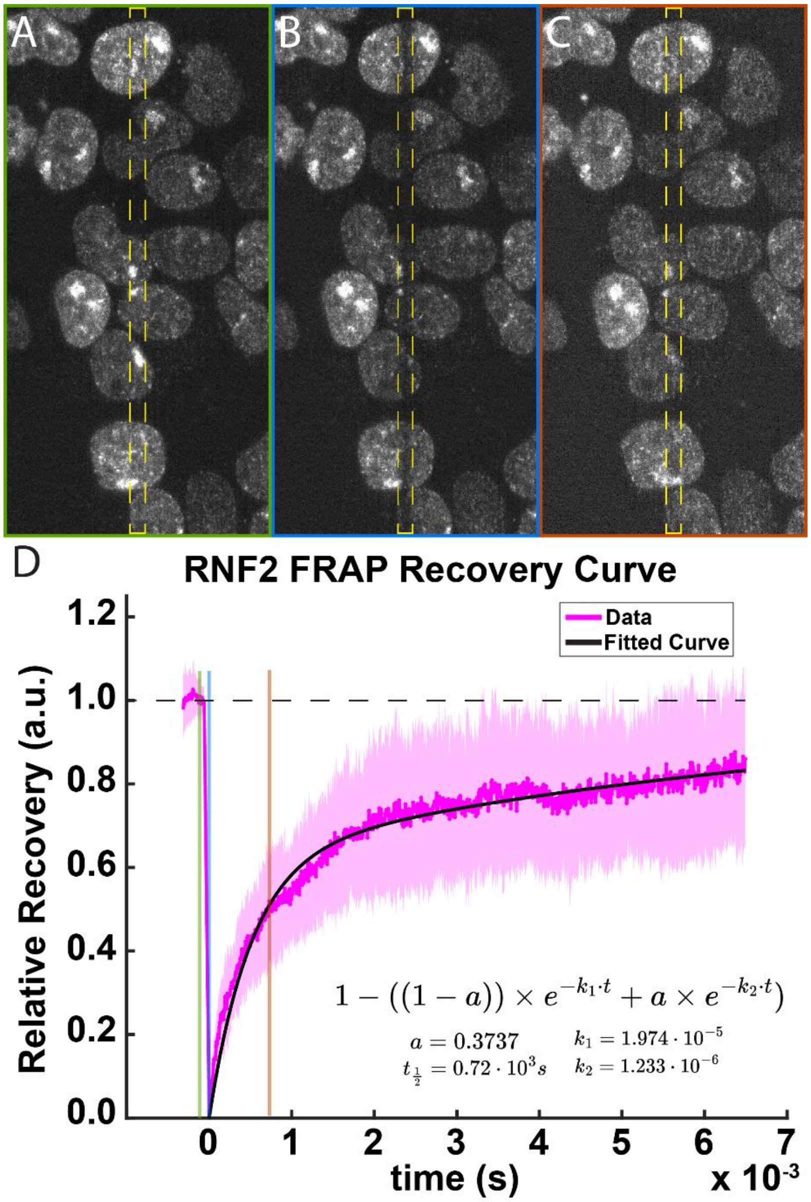

**Supplementary Figure 7.** FRAP analysis of RNF2 chromatin dynamics. **(A-C)** Representative confocal images of nuclei before photobleaching (A), immediately after photobleaching (B), and at the half-life for fluorescence recovery (C). **(D)** Normalized RNF2 FRAP recovery curve showing the experimental data (magenta line) with its standard deviation (magenta shaded region) and the fitted biexponential curve (black line). Recovery kinetics were fit using a two-component exponential model, yielding a fast and slow recovery component. The fitted mobile fraction parameter (a) and rate constants ($k_1$ and $k_2$) are indicated.

**Supplementary Table 1.** Bill of materials for the custom line-scanning confocal and HIST module with laser combiner and additional components for assembly

| Group/ Brand | Catalog Code | Parts | Count | Total Cost | Notes |
|---|---|---|---|---|---|
| Edmund Optics | EO-49359 | 85 x Ø25 mm VIS-NIR Achromat | 1 | $136.00 | |
| Kineflex | KINEFLEX-H-3-405..640-0.7-0.7-P (015064) | Qioptiq kineFLEX Fiber | 1 | $2,568.00 | *Quoted* |
| Kineflex | kineMATIX-P2 (011853) | Qioptiq kineMATIX Adapter/Mount | 2 | $1,194.00 | *Quoted* |
| Laserline Optics Canada | LOCP-8.9R10-0.8 | Powell Lens - 10deg Fan Angle, 0.8 mm Beam Width | 1 | $240.00 | |
| Laserline Optics Canada | LOCP-8.9R30-1.8 | Powell Lens - 30deg Fan Angle, 1.8 mm Beam Width | 1 | $240.00 | |
| McMasterCarr | 91292A004 | M2x4mm Socket Cap Screw | 4 | $14.76 | Pack of 100 |
| McMasterCarr | 91292A022 | M3x30mm Socket Cap Screw | 2 | $6.14 | Pack of 50 |
| McMasterCarr | 91292A107 | M4x6mm Socket Cap Screw | 2 | $9.09 | Pack of 100 |
| McMasterCarr | 91292A110 | M3x5mm Socket Cap Screw | 2 | $7.11 | Pack of 100 |
| McMasterCarr | 91292A113 | M3x10mm Socket Cap Screw | 8 | $7.11 | Pack of 100 |
| McMasterCarr | 91292A115 | M3x16mm Socket Cap Screw | 4 | $7.09 | Pack of 100 |
| McMasterCarr | 91292A116 | M4x10mm Socket Cap Screw | 5 | $10.34 | Pack of 100 |
| McMasterCarr | 91292A117 | M4x12mm Socket Cap Screw | 2 | $10.39 | Pack of 100 |
| McMasterCarr | 91292A123 | M3x20mm Socket Cap Screw | 2 | $9.69 | Pack of 100 |
| McMasterCarr | 91292A131 | M4x35mm Socket Cap Screw | 4 | $10.66 | Pack of 50 |
| McMasterCarr | 91292A137 | M6x20mm Socket Cap Screw | 6 | $6.88 | Pack of 25 |
| McMasterCarr | 91292A141 | M6x35mm Socket Cap Screw | 5 | $10.50 | Pack of 25 |
| McMasterCarr | 91292A441 | M6x10mm Socket Cap Screw | 14 | $9.54 | Pack of 50 |
| McMasterCarr | 91292A831 | M2x6mm Socket Cap Screw | 32 | $9.77 | Pack of 100 |

| | | | | | |
|---|---|---|---|---|---|
| McMasterCarr | 91292A832 | M2x8mm Socket Cap Screw | 6 | $7.74 | Pack of 100 |
| McMasterCarr | 91292A834 | M2x12mm Socket Cap Screw | 4 | $5.87 | Pack of 100 |
| McMasterCarr | 91585A189 | Dowel Pin Diameter 2mm Length 5mm | 35 | $17.53 | Pack of 100 |
| McMasterCarr | 91585A348 | Dowel Pin Diameter 3mm Length 8mm | 1 | $14.77 | Pack of 50 |
| McMasterCarr | 92235A105 | M4x8mm Flanged Alloy Socket Cap Screw | 6 | $16.96 | Pack of 50 |
| McMasterCarr | 92855A402 | M4x5mm Low Profile Socket Cap Screw | 1 | $3.93 | Pack of 10 |
| Minitec | - | Enclosure Box | 1 | $1,125.00 | *Quoted* |
| Misumi | CJPBB3-4 | SS Round Pin 3mm Shaft, 4mm Head | 7 | $96.95 | |
| Misumi | CJPDB3-4 | SS Diamond Pin 3mm Shaft, 4mm Head | 7 | $99.96 | |
| Nikon | - | 108 x Ø25 mm Lens | 1 | $593.45 | *Quoted* |
| OptoSigma | TADC-251C | Aluminum Linear X Stage 25mm Extended Contact Bearing Ways | 2 | $336.00 | |
| OptoSigma | TADC-251S | Aluminum Linear X Stage 25mm Extended Contact Bearing Ways | 2 | $336.00 | |
| OptoSigma | TADC-251SR | Aluminum Linear X Stage 25mm Extended Contact Bearing Ways | 3 | $504.00 | |
| Sciotex | MOSAIC-FPGA | FPGA | 1 | $7,940.00 | *Quoted* |
| SmarAct | MCS2-S-0005 | MCS2 Sensor Module in small aluminum housing | 1 | $940.00 | *Quoted* |
| SmarAct | MCS2-A-0600-150 | Sensor module cable with DSUB-15m connector and DSUB-15f connector | 1 | $140.00 | *Quoted* |
| SmarAct | MCS2-C-0007 | MCS2 control system in table top housing | 1 | $4,450.00 | *Quoted* |
| SmarAct | PWR-2C12D100-USA | Power Supply | 1 | $40.00 | *Quoted* |
| SmarAct | SLC-2445 | 45 mm Length Linear Piezo Stage | 1 | $2,500.00 | *Quoted* |
| SmarAct | SLC-2460 | 60 mm Length Linear Piezo Stage | 1 | $2,510.00 | *Quoted* |
| Thorlabs | AC254-125-A | 125mm x Ø1" Achromatic Doublet | 1 | $95.34 | |
| Thorlabs | AC300-050-A | 50mm x Ø30mm Achromatic Doublet | 1 | $108.04 | |
| Thorlabs | AC300-100-A | 100mm x Ø30mm Achromatic Doublet | 1 | $108.04 | |

| | | | | |
|---|---|---|---|---|
| Thorlabs | CLS-SL | Scan Lens with Large Field of View, 400 to 750 nm, EFL=70 mm | 1 | $3,262.86 |
| Thorlabs | GPS011-US | 1D or 2D Galvo System Linear Power Supply, 115 VAC | 1 | $582.95 |
| Thorlabs | GCE001 | Galvo Driver Card Cover | 1 | $69.00 |
| Thorlabs | GVS201 | 1D Galvo System, Broadband Mirror for 400-750 nm (-E02) | 1 | $1,662.38 |
| Thorlabs | LMRA9 | Ø0.5" Adapter for Ø9mm Optics | 2 | $38.90 |
| Thorlabs | MRA05-P01 | 5mm Right Angle Protected Silver Prism Mirror | 1 | $67.69 |
| Thorlabs | MRA15-P01 | 15mm Right Angle Protected Silver Prism Mirror | 1 | $89.61 |
| Thorlabs | MRAK25-P01 | 25 mm Knife-Edge Right-Angle Prism Prot. Silver Mirror | 1 | $156.35 |
| Thorlabs | PF05-03-P01 | Ø0.5" Protected Silver Mirror | 1 | $36.89 |
| Thorlabs | PF10-03-P01 | Ø1" Protected Silver Mirror | 2 | $120.40 |
| Thorlabs | PF20-03-P01 | Ø2" Protected Silver Mirror | 2 | $245.50 |
| Thorlabs | POLARIS-K05 | Ø1/2" Mirror Mount, 3 Low-Profile Hex Adjusters | 3 | $530.04 |
| Thorlabs | POLARIS-K2S3 | Ø2" Mirror Mount, 3 Low-Profile Hex Adjusters | 2 | $648.24 |
| Thorlabs | PRM05-M | Ø0.5" High-Precision Rotation Mount | 2 | $416.94 |
| Thorlabs | SM05L03 | SM05 Lens Tube, 0.3" Depth | 1 | $15.96 |
| Thorlabs | SM05L10 | SM05 Lens Tube, 1" Depth | 2 | $34.94 |
| Thorlabs | SM1L10 | SM1 Lens Tube, 1" Depth | 1 | $16.49 |
| Thorlabs | SM1L15 | SM1 Lens Tube, 1.5" Depth | 1 | $18.17 |
| Thorlabs | SM1L20 | SM1 Lens Tube, 2" Depth | 1 | $19.10 |
| Thorlabs | SM1L25 | SM1 Lens Tube, 2.5" Depth | 1 | $24.43 |
| Thorlabs | SM30L05 | SM30 Lens Tube, 0.5" Depth | 1 | $34.96 |
| Thorlabs | SM30L10 | SM30 Lens Tube, 1" Depth | 1 | $37.50 |
| Thorlabs | SM30L30 | SM30 Lens Tube, 2" Depth | 1 | $48.31 |
| Thorlabs | TTL200MP | Laser Scanning Tube Lens, f = 200 mm | 1 | $1,627.28 |
| Thorlabs | VA100-M | Adjustable Mechanical Slit, M4 Tap, Metric Micrometer | 1 | $321.57 |

| | | | | | |
|---|---|---|---|---|---|
| Custom Parts | - | 1" Mirror Mount to 0.5" Polaris | 1 | $20.37 | *JLCCNC* |
| Custom Parts | - | 30mm Lens Tube Mount | 1 | $38.14 | *JLCCNC* |
| Custom Parts | - | Base Plate | 1 | $261.86 | *JLCCNC* |
| Custom Parts | - | Extension Plate | 1 | $240.46 | *JLCCNC* |
| Custom Parts | - | Galvo Mount | 1 | $23.22 | *JLCCNC* |
| Custom Parts | - | Kineflex and Split Path Mirror Mount | 1 | $53.01 | *JLCCNC* |
| Custom Parts | - | Knife Edge Mirror Mount for Manual Stage | 1 | $15.97 | *JLCCNC* |
| Custom Parts | - | Knife Edge Mirror Mount for SmarAct Stage | 1 | $15.89 | *JLCCNC* |
| Custom Parts | - | Mirror to Manual Stage Adapter | 1 | $12.53 | *JLCCNC* |
| Custom Parts | - | Mirror to SmarAct Stage Adapter | 1 | $15.35 | *JLCCNC* |
| Custom Parts | - | Mounting Port to Base Plate Adapter | 1 | $51.54 | *JLCCNC* |
| Custom Parts | - | Polaris on Stage Mount | 1 | $12.56 | *JLCCNC* |
| Custom Parts | - | Rotation Mount to Stage Adapter | 2 | $30.98 | *JLCCNC* |
| Custom Parts | | SM05 Lens Tube Mount | 1 | $16.86 | *JLCCNC* |
| Custom Parts | - | SM1 Lens Tube Mount | 1 | $33.88 | *JLCCNC* |
| Custom Parts | - | SM2 Lens Tube Mount | 2 | $52.08 | *JLCCNC* |
| Custom Parts | - | Stage to Stage Adapter | 2 | $44.44 | *JLCCNC* |
| Custom Parts | - | Support Beam 1 (for Extension Plate) | 1 | $20.12 | *JLCCNC* |
| Custom Parts | - | Support Beam 2 (for Extension Plate) | 1 | $13.47 | *JLCCNC* |
| Custom Parts | - | Support Pillar (for Extension Plate) | 2 | $20.66 | *JLCCNC* |
| Custom Parts | - | Variable Slit Mount | 1 | $12.73 | *JLCCNC* |
| | | | **Subtotal** | $37,629.23 | |

| ***Laser Combiner*** | | | | | |
|---|---|---|---|---|---|
| AA Optoelectronics | AOTFnC-400650-CPCH-AS-PRO-257 | AO Tunable Filter | 1 | $2,444.40 | *Quoted* |
| AA Optoelectronics | MPDS4C-B66-22-52.111-RS-BT0 | Multi-Purpose Digital Synthesizer 4 channels | 1 | $2,388.00 | *Quoted* |
| Bolder Vision Optik | AHWP3 | Ø0.5" VIS Half Wave Plate | 1 | $395.00 | *Quoted* |
| Edmund Optics | EO-68329 | Ø25 mm Silver Mirror | 1 | $146.00 | |

| | | | | | |
|---|---|---|---|---|---|
| McMasterCarr | 91292A004 | M2x4mm Socket Cap Screw | 16 | $14.76 | Pack of 100 |
| McMasterCarr | 91292A038 | M4x14mm Socket Cap Screw | 14 | $13.57 | Pack of 100 |
| McMasterCarr | 91292A113 | M3x10mm Socket Cap Screw | 2 | $7.11 | Pack of 100 |
| McMasterCarr | 91292A116 | M4x10mm Socket Cap Screw | 1 | $10.34 | Pack of 100 |
| McMasterCarr | 91292A121 | M4x20mm Socket Cap Screw | 2 | $12.98 | Pack of 100 |
| McMasterCarr | 91292A123 | M3x20mm Socket Cap Screw | 2 | $9.69 | Pack of 100 |
| McMasterCarr | 91292A832 | M2x8mm Socket Cap Screw | 20 | $7.74 | Pack of 100 |
| McMasterCarr | 91585A189 | Dowel Pin Diameter 2mm Length 5mm | 14 | $17.53 | Pack of 100 |
| McMasterCarr | 91585A271 | Dowel Pin Diameter 2.5mm Length 6mm | 1 | $10.25 | Pack of 50 |
| MPB | 2RU-VFL-P-500-488-B1R | Blue Visible Fiber Lasers | 1 | $20,947.50 | *Quoted* |
| MPB | 2RU-VFL-P-2000-642-B1R | Red Visible Fiber Lasers | 1 | $13,205.00 | *Quoted* |
| MPB | 2RU-VFL-P-1000-560-B1R | Green Visible Fiber Lasers | 1 | $17,910.00 | *Quoted* |
| OptoSigma | TADC-251S | Aluminum Linear X Stage 25mm Extended Contact Bearing Ways | 1 | $168.00 | |
| OptoSigma | TADC-251SR | Aluminum Linear X Stage 25mm Extended Contact Bearing Ways | 4 | $672.00 | |
| Semrock | LM01-427-25 | Dichroic Mirror | 1 | $289.28 | |
| Semrock | LM01-503-25 | Dichroic Mirror | 1 | $289.28 | |
| Semrock | LM01-613-25 | Dichroic Mirror | 1 | $289.28 | |
| Thorlabs | BB05-E02 | Ø1/2" Broadband Dielectric Mirror, 400 - 750 nm | 2 | $119.22 | |
| Thorlabs | GBE02-B | 2X Achromatic Galilean Beam Expander, AR Coated: 650 - 1050 nm | 1 | $512.24 | |
| Thorlabs | LB1 | Beam Block, 400 nm - 2 µm, 10 W Max Avg. Power | 1 | $62.92 | |
| Thorlabs | MRA10-EO2 | Right-Angle Prism Dielectric Mirror, 400 - 750 nm, L = 10.0 mm | 4 | $491.20 | |

| | | | | | |
|---|---|---|---|---|---|
| Thorlabs | PBS101 | 10 mm Polarizing Beamsplitter Cube, 420 - 680 nm | 2 | $475.38 | |
| Thorlabs | POLARIS-K05S1 | Ø1/2" Mirror Mount, 2 Low-Profile Hex Adjusters | 1 | $163.97 | |
| Thorlabs | POLARIS-K1S5 | Ø1" Mirror Mount, 3 Hex Adjusters with Side Holes, Monolithic Optic Retention | 4 | $963.60 | |
| Thorlabs | RSP05/M | Rotation Mount for Ø1/2" (Ø12.7 mm) Optics, M4 Tap | 1 | $89.72 | |
| Toptica | Toptica IBEAM-SMART-405-S-HP | Single Mode Diode Laser 405 nm | 1 | $6,985.00 | *Quoted* |
| Custom Parts | - | Laser Combiner Baseplate | 1 | $139.52 | *JLCCNC* |
| Custom Parts | - | Prism-to-X-Stage Mounting Plate | 4 | $51.16 | *JLCCNC* |
| Custom Parts | - | Beam Steering Assembly Backplate | 4 | $109.84 | *JLCCNC* |
| Custom Parts | - | HWP Mounting Plate | 1 | $24.74 | *JLCCNC* |
| Custom Parts | - | Polarizing Beamsplitter Mounting Block | 1 | $25.09 | *JLCCNC* |
| Custom Parts | - | Small Mirror to Translation Stage Adapter | 1 | $18.80 | *JLCCNC* |
| Custom Parts | - | AOTF Beam Dump | 1 | $15.36 | *JLCCNC* |
| Custom Parts | - | AOTF Mount | 1 | $41.39 | *JLCCNC* |
| Custom Parts | - | Beam Expander Mount | 1 | $20.61 | *JLCCNC* |
| | | | **Subtotal** | $69,557.47 | |

| | | | | | |
|---|---|---|---|---|---|
| ***Nikon Accessories*** | | | | | |
| - | - | 135 x Ø25 mm Internal Nikon Lens | 1 | $0.00 | |
| Chroma | | Dichroic Mirror (for Nikon) | 1 | $1,225.00 | *Quoted* |
| Hamamatsu/Nikon | ORCA-FUSION | ORCA Fusion Camera | 1 | $25,988.18 | *Quoted* |
| Nikon | CFI-SR-Plan-Apo-IR-60x | CFI-SR-Plan Apo-IR - 60x 1.27 NA Water Immersion Objective | 1 | $21,532.29 | *Quoted* |
| Nikon | QGNPS083-NI | Queensgate Piezo Stage | 1 | $12,703.80 | *Quoted* |
| | | | **Subtotal** | $61,449.27 | |

| | | | | |
|---|---|---|---|---|
| ***TIRF Components*** | | | | |
| Thorlabs | AC080-010-A | 10mm x Ø8mm Achromatic Doublet | 1 | $58.81 |

| | | | | |
|---|---|---|---|---|
| Thorlabs | AC254-200-A | 200mm x Ø1" Achromatic Doublet | 1 | $95.34 |
| Thorlabs | AD8T | Ø1inch OD Adapter for Ø8 mm Optic | 1 | $22.75 |
| Thorlabs | SM1L30 | SM1 Lens Tube, 3" Depth | 1 | $31.28 |
| Thorlabs | SM1L35 | SM1 Lens Tube, 3.5" Depth | 1 | $43.38 |
| | | | **Subtotal** | $251.56 |

| | |
|---|---|
| **TOTAL** | $131,258.30 |

**Supplementary Table 2.** Comparison of effective FOV for each objective and mode between the design specification and experimental measurements.

| Objective | Mode | Design FOV | Experimental FOV |
|---|---|---|---|
| 40x | *HIST* | 375 x 375 µm | 375 x 345 µm |
| | *Line-scanning Confocal* | 375 x 375 µm | 375 x 225 µm |
| 60x | *HIST* | 250 x 250 µm | 250 x 227.5 µm |
| | *Line-scanning Confocal* | 250 x 250 µm | 250 x 160 µm |
| 100x | *HIST* | 150 x 150 µm | 150 x 115 µm |
| | *Line-scanning Confocal* | 150 x 150 µm | 150 x 90 µm |

**Supplementary Table 3.** Detailed imaging parameters.

| Figure | Excitation Mode | Objective (NA) | Wavelength (nm) | Exposure/Cycle time (ms) |
|---|---|---|---|---|
| 2A | Confocal | 60x (1.27) | 560 | 100.0/130.0 |
| 2B | HIST | 60x (1.27) | 560 | 100.0/130.0 |
| 3A | Widefield | 60x (1.27) | 560 | 50.0/50.7** |
| 3B | HIST | 60x (1.27) | 560 | 50.0/65.0 |
| 3C | Confocal | 60x (1.27) | 560 | 50.0/65.0 |
| 3D | Confocal | 60x (1.27) | 560 | 50.0/65.0 |
| 4A, B | Confocal | 60x (1.27) | 405 | 100.1/125.0 |
| | | | 560 | 100.0/125.0 |
| | | | 642 | 100.0/125.0 |
| 4C-E | HIST | 60x (1.27) | 560 | 100.3/125.0 |
| | | | 642 | 100.0/111.3 |
| 5A | Confocal | 60x (1.27) | 488 | 100.0/130.0 |
| | HIST | | 642 | 20.0/25.0 |
| S2A | Confocal | 40x (1.25) | 560 | 100.0/130.0 |
| S2B | HIST | 40x (1.25) | 560 | 100.0/130.0 |
| S3A | Confocal | 100x (1.35) | 560 | 50.0/63.2 |
| S3B | HIST | 100x (1.35) | 560 | 50.0/63.2 |

| | | | | |
|---|---|---|---|---|
| S4A | Widefield | 40x (1.25) | 560 | 50.0/50.7** |
| S4B | HIST | 40x (1.25) | 560 | 50.0/65.0 |
| S4C | Confocal | 40x (1.25) | 560 | 50.0/65.0 |
| S4D | Confocal | 40x (1.25) | 560 | 50.0/65.0 |
| S5A | Widefield | 100x (1.35) | 560 | 50.0/50.7** |
| S5B | HIST | 100x (1.35) | 560 | 50.0/65.0 |
| S5C | Confocal | 100x (1.35) | 560 | 50.0/65.0 |
| S5D | Confocal | 100x (1.35) | 560 | 50.0/65.0 |
| S6B | Widefield | 60x (1.49) | 560 | 50.0/61.3** |
| S6C | TIRF | 60x (1.49) | 560 | 50.0/61.3** |
| S7 | Confocal | 60x (1.27) | 642 | 25.0/325.0 |

** Lightsheet readout mode was disabled
